# Infestation dynamics between parasitic Antarctic fish leeches (Piscicolidae) and their crocodile icefish hosts (Channichthyidae)

**DOI:** 10.1101/2020.01.07.897496

**Authors:** Elyse Parker, Christopher Jones, Patricio M. Arana, Nicolás A. Alegría, Roberto Sarralde, Francisco Gallardo, A.J. Phillips, B.W. Williams, A. Dornburg

## Abstract

An understanding of host-parasite interactions represents an important, but often overlooked, axis for predicting how marine biodiversity may be impacted by continued environmental change over the next century. For host and parasite communities in the Southern Ocean, investigations of many major groups of parasites have largely been limited to taxonomic and phylogenetic studies, creating an urgent need for the collection of baseline ecological data if we are to detect changes in host-parasite interactions in the future. Here, we survey three species of Crocodile icefish (Notothenioidei: Channichthyidae) collected from two island archipelagos in Antarctica’s South Scotia Arc region for evidence of leech infestations. Specifically, we report on infestation prevalence and intensity of three leech species (*Trulliobdella bacilliformis*, *Trulliobdella capitis*, and *Nototheniobdella sawyeri*) on the host fish species *Chaenocephalus aceratus*, *Champsocephalus gunnari*, and *Chionodraco rastrospinosus.* Additionally, we characterize spatial patterns of relative abundances of each leech species across the Elephant and South Orkney Islands, size distribution of parasitized fish, and patterns of host and attachment site specificity. Our results reveal high levels of attachment area fidelity for each leech species. These results suggest skin thickness and density of the vascular network constrain leech attachment sites and further suggest trophic transmission to be an important axis of parasitization. We also demonstrate that, while leech species appear to be clustered spatially, this clustering does not appear to be correlated with fish biomass. This study illuminates the complex interactions among fish hosts and leech parasites in the Southern Ocean and lays the groundwork for future studies of Antarctic marine leech ecology that can aid in forecasting how host-parasite interactions may shift in the face of ongoing climate change.

## Introduction

Over the past several decades, Antarctica’s Southern Ocean has become increasingly threatened by climate change, fisheries interests, and a general increase in human traffic (Kennicutt et al. 2015). The impact of these changes on the future conservation status of wildlife is an area of intense research interest (Kock 2007; Raymond et al. 2014; Gutt et al. 2015; Krüger et al. 2017). However, lack of baseline data for Antarctica’s parasite communities has challenged our ability to integrate these organisms in forecasts of Southern Ocean biodiversity over the next century (Bielecki et al. 2008). Differential responses to environmental stressors by hosts and their parasites can alter the balance of host-parasite interactions (Gehman et al. 2018), creating an urgent need for the quantification of baseline infestation parameters of ecologically or economically important host species (Palm 2011). Unfortunately, our knowledge of the ecology and dynamics of infestation for many groups of parasites in the Southern Ocean are restricted to few studies, and many aspects of their biology remain virtually unexplored.

Across the world, marine fish leech (Piscicolidae) species are major pests for commercially important fisheries (Cruz-Lacierda et al. 2000; Marancik et al. 2012) and also epidemiologically important as vectors transmitting viruses and additional blood parasites (Khan 1980, 1984; Siddall and Desser 1993; Karlsbakk 2004). Although the few studies reporting leech abundances on fishes from the Southern Ocean have indicated a general low prevalence, leeches are spatially heterogeneous with high loads on individual fish (Bielecki et al. 2008; Kuhn et al. 2018). This distributional pattern is common for parasitic organisms in general (May 1978; Pacala et al. 1990) and suggests it is likely that Antarctica’s fishes face parasitic pressures similar to those of fishes in other parts of the world (Sawyer and Hammond 1973). However, the Southern Ocean is unusual in that the vast majority of marine teleost fish biomass, species abundance, and diversity is dominated by one clade: notothenioids (Eastman 2000). These fishes include species of high economic importance to fisheries (Kock 1992; Delord et al. 2009; Collins et al. 2010) and species that represent critical links in the Antarctic food web between lower trophic levels and higher level consumers such as whales, seals, and penguins (Targett 1981; Smith et al. 2007). However, the impact of piscicolids on notothenioids remains unclear. As future ecological relationships of the Southern Ocean become increasingly uncertain, developing a basic understanding of marine leeches now is critical if we are to be able to detect changes in host-parasite interactions or disease transmission pathways in the future. How prevalent are leech infestations? Do leeches disproportionately parasitize one size class over another? Do sympatric species of leeches compete for the same host fish? Answering fundamental questions such as these is critical if we are to illuminate the role of leeches in the current and future ecology of the Southern Ocean.

Crocodile icefishes (Channichthyidae) are an exemplar lineage from which to investigate the dynamics of leech infestation in notothenioids. Globally renowned for their lack of hemoglobin (Sidell and O’Brien 2006), these scaleless fishes are among the most well-studied teleosts in polar latitudes (Ruud 1954; Cocca et al. 1995; Kock 2005; Near et al. 2012; Melillo et al. 2015; Giraldo et al. 2016), and contain species such as mackerel icefish (*Champsocephalus gunnari*) that are of high economic importance for fisheries (Williams et al. 1994; Duhamel et al. 1995; Everson et al. 1999; Everson 2015). Correspondingly, this is one of few clades in which leech occurrences and identifications are well-documented by species (Bielecki et al. 2008; Oguz et al. 2012; Kuhn et al. 2018), providing the necessary taxonomic framework for ecological study

Prior investigations of Antarctic leech parasitism have suggested three species to be particularly common across a range of channichthyid host species: *Nototheniobdella sawyeri* A. Utevsky, 1993, *Trulliobdella bacilliformis* (Brinkmann,1947), and *Trulliobdella capitis* Brinkmann, 1947. *Trulliobdella capitis* and *T. bacilliformis* were described from the dorsal region of the head of the blackfin icefish, *Chaenocephalus aceratus*, and from the oral cavity of the South Georgia icefish, *Pseudochaenichthys georgianus*, respectively (Brinkmann 1948). Since then, little data has been collected on the ecology of *T. bacilliformis*. While this species has also been discovered to infest at least four other species of icefish, prevalence and infection intensity of this leech have not been thoroughly investigated (Table 1). In contrast, *T. capitis* has been identified from eight additional channichthyid species, with patterns of infestation by *T. capitis* examined for five of these species. These studies suggest a generally low prevalence (3-13%) across host species with a range of attachment sites including the head, body, mouth, and gills (Table 1). A notable exception for this pattern stems from Bielecki et al. (2008) who reported a prevalence of 45% in *Chionodraco rastrospinosus* with attachment sites concentrated in the cranial areas of the species (Table 1). However, absence of additional data from this host-parasite species pair challenges interpretation of whether this high prevalence is a general condition or a localized concentration of parasite activity.

**Table 1:**
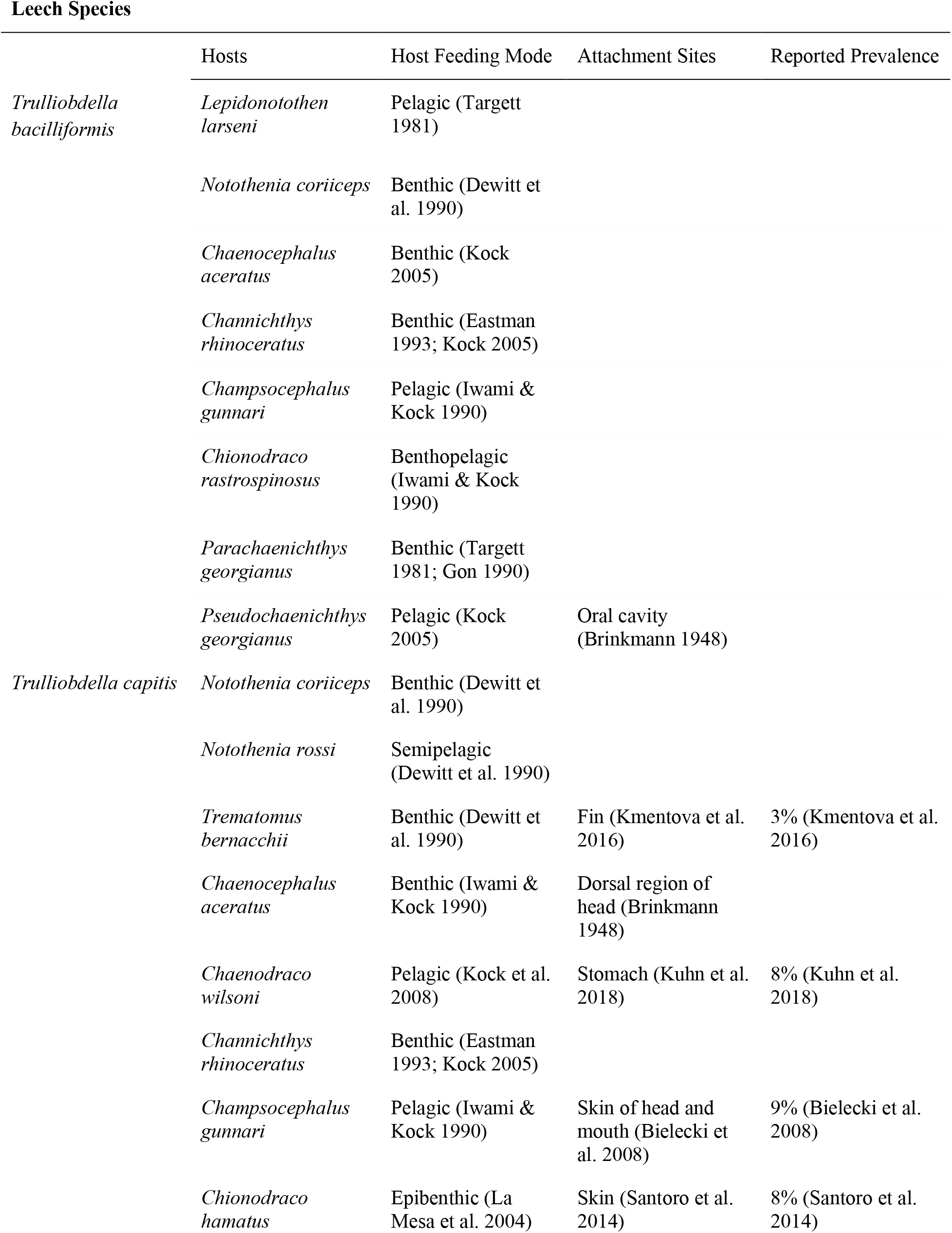
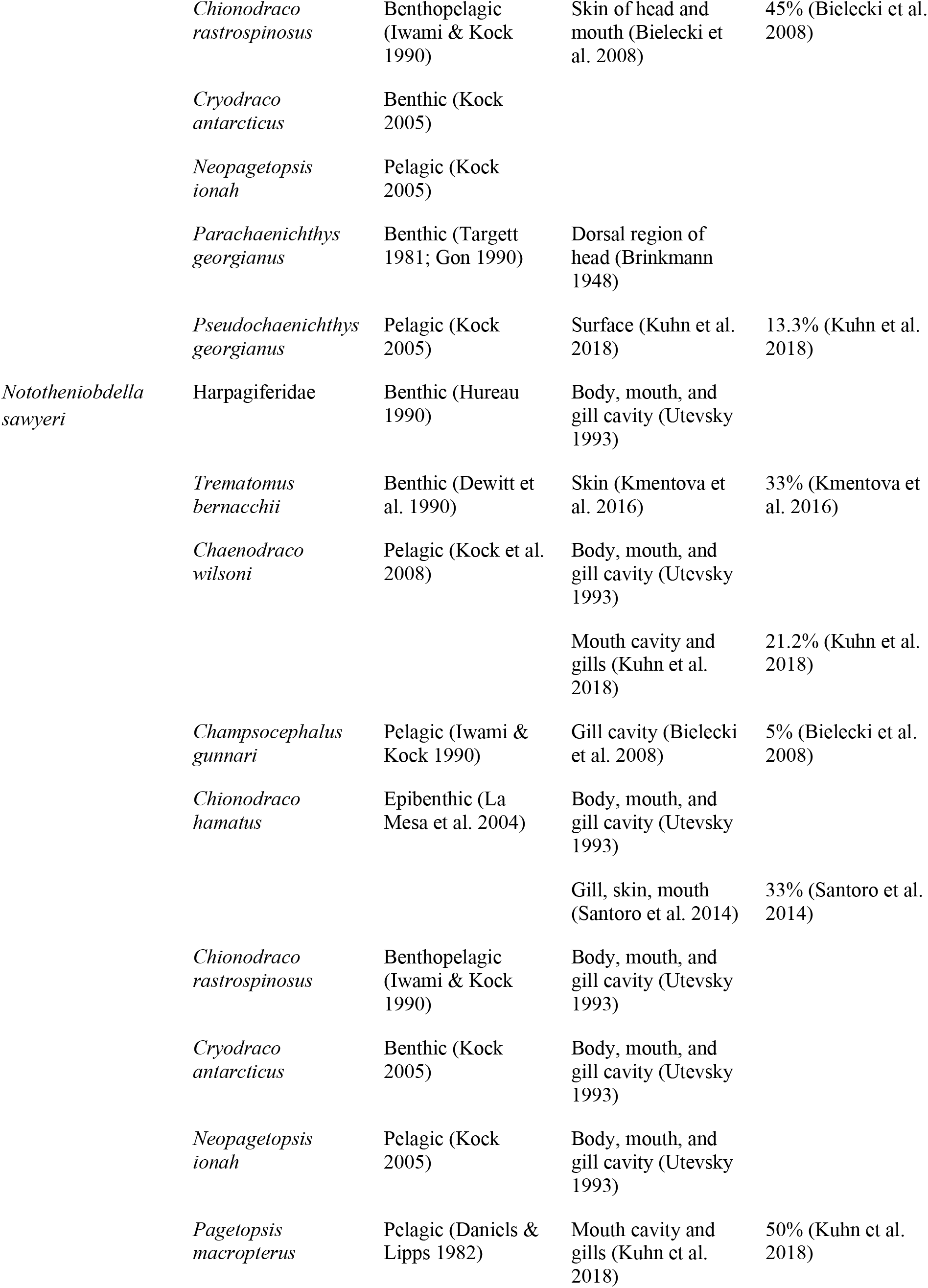
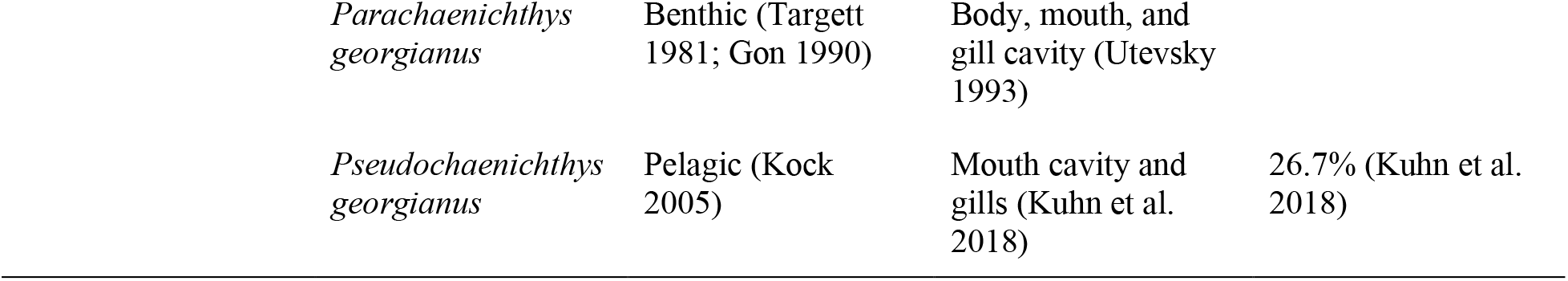
Summary of reported infection parameters for three Antarctic leech species and each of their notothenioid fish hosts.

Relative to either species of *Trulliobdella, N. sawyeri* has received more study since its initial description (Utevsky 1993). *Nototheniobdella sawyeri* has been reported from the body surface, mouth, and gill cavity of eight channichthyid species (Utevsky 2005; Bielecki et al 2008; Kuhn et al. 2018), including infestation parameters for at least five channichthyid species (Table 1). Further contrasting observations of *T. capitis* with *N. sawyeri* suggests the latter to generally have higher prevalence (20-50%) than the former (8-13%) across the range of channichthyid hosts (Table 1). However, the commercially harvested mackerel icefish, *Champsocephalus gunnari*, diverges from this pattern with a prevalence of only 5% (Bielecki 2008). Lack of further investigation again stymies interpretation of whether this finding of lower prevalence is anomalous or represents a general trend for this host species. More broadly, these knowledge gaps are unfortunate as channichthyids are widespread around the entire Antarctic, often occurring at extremely high biomass (Reid et al. 2007), with individual fishes harboring high infestation intensities of leeches (Bielecki et al. 2008; Kuhn et al. 2018). The limited number of detailed investigations of infestation parameters, not only creates a barrier to our understanding of leech-host ecology in the Antarctic, but also challenges our ability to establish general trends of infestation required for monitoring host fish stock health.

The purpose of this study was to characterize patterns of leech infestation on notothenioid fishes collected from two island archipelagos in Antarctica’s South Scotia Arc. Here we report on various parameters related to leech infestations of three species of Crocodile icefish (Notothenioidei: Channichthyidae) distributed around Elephant Island and the South Orkney Islands. Investigated infestation parameters included: (1) the proportion of each leech species found to parasitize Crocodile icefish taxa; (2) the relative frequencies of each leech species at each collection site across the Elephant and South Orkney Islands; (3) the size distribution of parasitized fish relative to the size distribution of all fish collected; and (4) the prevalence and intensity of each leech species on each crocodile icefish species. We assess both global trends as well as differences in infestation parameters between Elephant Island and the South Orkney Islands, thereby revealing previously undocumented aspects of marine leech ecology in the Southern Ocean that include infestation site fidelity between host-parasite species pairs and host size preferences.

## Materials and Methods

### Sample acquisition

Sampling was conducted aboard the R/V Cabo de Hornos (Call Sign = CB7960) between 6–27 January, 2018, as part of the Chilean finfish bottom trawl survey (Delegation of Chile, 2018) of finfish around Elephant Island and the South Orkney Islands in the Southern Ocean. Trawling was conducted using a Hard Bottom Snapper Trawl (HBST; Net Systems, Inc., Bainbridge Island, WA) and a Cassanova trawl. Thirty-five sampling sites were selected based on a random, depth stratified survey design that was primarily restricted to shelf regions (50–500 m). During trawl deployment, sensors monitored the geometry and seabed bottom contact of the trawl, ensuring that the trawl ground tackle was in contact with the seafloor for a 30-minute tow.

All collected fish were surveyed for evidence of past and present leech infestation, yielding a total of 89 individual parasitized fishes representing three species and genera of Crocodile icefishes (Channichthyidae). Locations of leech infestations were recorded and photographed for each parasitized fish. Identification and counts of leech specimens were conducted during the survey, and voucher photographs and specimens of both leeches and their host fish, were deposited in the North Carolina Museum of Natural Sciences (Supplemental materials). The total length and weights of all fishes captured were additionally recorded to enable quantification of leech abundances as they relate to fish size.

### Leech infestation parameters

Several general parameters were examined to assess overall trends of leech infestation, with terminology following Bush et al. (1997). First, we calculated the relative proportions of each leech species that were found to parasitize each species of icefish in order to investigate patterns of host-parasite specificity. We also calculated the relative proportions of each leech species collected from each sampling site across the Elephant and South Orkney islands. Parasite infestation intensities per individual were quantified and compared to the distribution of total fish caught (i.e. prevalence). Additionally, we explicitly tested the effect of body size on leech infestation using ANOVA in combination with a Tukey honest significance test. This allowed us to compare the distribution of total lengths of all parasitized fish to the distribution of total lengths of all =non-parasitized fish to investigate whether there exists a correlation between fish size class and frequency of parasitism. Tests were conducted for each icefish species, both across islands and for Elephant Island and the South Orkney Islands individually.

We quantitatively assessed differences in leech occurrences between fish species by generating violin plots using the vioplot package in R (Adler 2005). These plots modify a traditional box plot by adding a rotated kernel density plot, thereby facilitating inspection of quartiles and the underlying probability distribution of the data (Hintze and Nelson 1998). We mapped distribution of leeches to specific body regions of the host, dividing the body into seven areas that included all areas leeches were encountered: dorsal surface of the head; lateral surfaces of the head; ventral surface of the head; inside of upper jaw; inside of lower jaw; ventral region of the body; and the pelvic fins. Although some Antarctic leech species are described as occurring on the gills of Antarctic fishes, no leeches represented in this study were found on gills and therefore this region was not included in our division of host bodies.

To account for the potential impact of small sample sizes on the qualitative interpretation of patterns related to leech infestation, a number of simulations were performed to test whether measures of infestation differed significantly from what might be expected based on random sampling alone. We investigated whether host fish species predicts relative proportion of leech species observation. For this test, we simulated a dataset of 1,000 individuals for each species of host fish, and each individual was randomly assigned to host one of the three species of leech. From this simulated dataset, we randomly sampled a subset of individuals, with subsample size mirroring the empirical sample sizes for each fish species in this study. Sampling was repeated 5,000 times for each subsample size. For each host fish species, we then compared the simulated distribution of each leech species’ percent frequencies to the empirically calculated relative frequencies of each leech species observed on each fish species to assess significance.

We also investigated whether particular leech species preferentially infest specific body regions on their host fish. To evaluate this, the host body was divided into discrete areas on which leeches were observed. We simulated a dataset of equal numbers (1,000) of each host fish species collected for each of two focal leech species: 1) *T. capitis* and 2) *T. bacilliformis*. To each of the individuals from each host fish species, we randomly assigned one of the seven body regions in which a leech was observed. From this simulated dataset, we randomly sampled a subset of individuals, with subsample size mirroring the empirical sample sizes for each fish species in this study. Sampling was repeated 5,000 times for each subsample size. For each of the two focal leech species, we then compared the simulated distributions of infestations across body regions to the empirically calculated relative proportion of infestation to assess significance. All simulations were conducted in R.

### Comparing estimates of fish and leech biomass

We tested whether leech abundance was correlated with fish abundance or biomass using two approaches. First, we tested if fish abundance predicted leech abundances in our samples using a simple linear regression model. We used the “linear model” (lm) function in R to calculate the correlation between the number of parasitized fish and the total number of fish collected at each station across both Elephant Island and the South Orkney Islands. This analysis was performed independently for each host fish species. Second, we estimated total standing biomass for each host fish and *T. bacilliformis* and *T. capitis* over the total shelf area (50 m–500 m) of Elephant Island and the South Orkney Islands. For each haul, total count of individual leech species captured for all finfish combined was summed and standardized to one square nautical mile of area swept using the average trawl mouth width and bottom distance covered at each station. Estimates of standing stock biomass were computed using a Delta-lognormal maximum likelihood approach (Pennington 1985; De la Mare 1994), with overall biomass *B* as

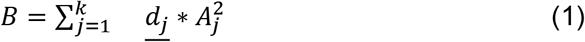

where the biomass is a product of the mean density *d*_*j*_ of the sampling strata *j* and the area *A*_*j*_ of that strata). Similarly, variance 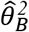 of abundance estimates are a product of the strata area squared, where

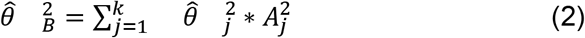

Seabed areas for the Elephant Island shelf (50-500 m) were taken from Jones et al. (1999) and for the South Orkney Islands from Jones (2000). The above equations were again used to estimate fish biomass. This approach allowed us to both visualize areas of high leech or fish biomass, and incorporate trawl parameters into expectations of biomass when repeating the statistical tests outlined above.

## Results

### Patterns of host and site specificity

Out of all 1,461 fish examined, we encountered 89 individuals with leech infestations, yielding a combined prevalence of 6.1 %. Across both Elephant Island and the South Orkney Islands, we encountered 426 leeches on a total of 89 individual fish: 30 individuals of *C. aceratus*, 43 individuals of *C. gunnari*, and 16 individuals of *C. rastrospinosus*. Based on this pooled sampling of individuals from the two regions, we calculated the relative proportion of infestation (RPI) of all three leech species (*T. capitis*, *T. bacilliformis*, and *N. sawyeri*) that parasitized each host fish species (Table 2). These proportions are suggestive of patterns of a degree of host-parasite specificity across the Elephant and South Orkney islands, with *T. bacilliformis* appearing to preferentially infest *C. gunnari* (80% RPI; Table 2) and *T. capitis* appearing to preferentially infest *C. aceratus* (54% RPI; Table 2). The results of our simulations demonstrate that for *C. aceratus*, the RPI by *T. capitis* is higher than would be expected under an assumption of random assortment (ESM Fig. 1; RPI = 0.54, p = 0.0084). The same is true of the empirical RPI of *C. gunnari* by *T. bacilliformis* (ESM Fig. 1, RPI = 0.71, p=0.000), and the empirical RPI of *C. rastrospinosus* by *N. sawyeri* (ESM Fig. 1; RFI = 0.8, p=0.000).

**Table 2:** Relative proportion of infestation of each leech species per host fish species across pooled samples from Elephant Island and the South Orkney Islands. Greatest relative proportion of each leech species is bolded.

|  | Relative proportion of *Trulliobdella bacilliformis* | Relative proportion of *Trulliobdella capitis* | Relative proportion of *Nototheniobdella sawyeri* |
| --- | --- | --- | --- |
| *Chaenocephalus aceratus* | 22% | **54%** | 0% |
| *Champscephalus gunnari* | **71%** | 24% | 20% |
| *Chionodraco rastrorpinosus* | 7% | 22% | **80%** |

**Figure 1:**
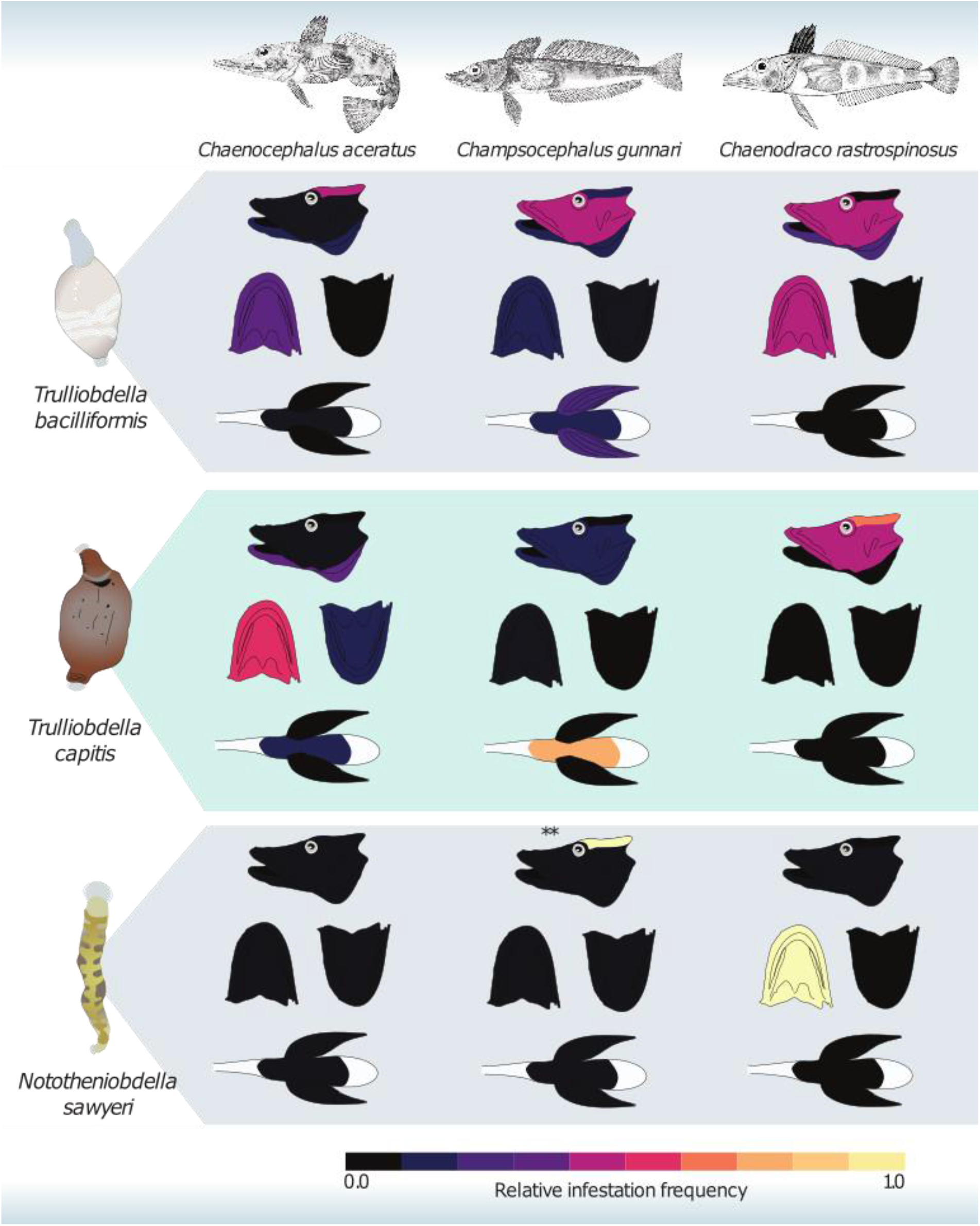
Relative infestation frequency of the Antarctic fish leech species *Trulliobdella bacilliformis* (top); *Trulliobdella capitis* (middle); and *Nototheniobdella sawyeri* (bottom) on three species of Crocodile icefishes. Brightness indicates relative frequency of infestation on body surfaces (dorsal, lateral, and ventral surfaces of the head; upper jaw; lower jaw; ventral surface of body between pelvic fins; and pelvic fins). ** indicates single *N. sawyeri* found inside the braincase of a single *Champsocephalus gunnari*. Fish illustrations modified from public domain images with Creative Commons Licenses.

Quantifying the distribution of leeches on host fish additionally suggests site specificity of leech species on particular host fish species (Fig. 1). Our results demonstrate that the species *T. capitis* most frequently infests *C. aceratus* on the upper inner jaw [Relative infestation frequency (RFI) = 0.52; Fig. 1], while on *C. gunnari* it is most commonly encountered between the pectoral fins (RFI = 0.8333; Fig. 1) and on *C. rastrospinosus*, occurs most frequently on the dorsal surface of the head (RFI = 0.6087;Fig. 1). *T. bacilliformis* appears to preferentially infest the dorsal surface of the head on *C. aceratus* (RFI = 0.667; Fig. 1), but on *C. gunnari* it is found predominantly on the lateral surfaces of the head (RFI = 0.3898; Fig. 1). Simulation results demonstrate that for both *T. capitis* and *T. bacilliformis,* these empirical infestation frequencies on particular body regions of host fish species are higher than the frequencies that might be expected under an assumption of random assortment (Fig. 1 & ESM Fig. 1).

There were only seven observations of *T. bacilliformis* on *C. rastrospinosus*, with 2 observations each on the lateral surfaces of the head, on the inner region of the lower jaw, and on the inner region of the upper jaw, but the empirical RFI values of these infestation parameters did not differ significantly from expectations of random assortment (ESM Fig. 1). From our sample, we also observed 6 individuals of *C. gunnari* and 2 individuals of *C. aceratus* from the South Orkney Islands that were parasitized by both *T. capitis* and *T. bacilliformis* simultaneously (ESM Table 1). On two individuals of *C. gunnari*, members of the two leech species were found together in the same body region (between the pelvic fins on one individual and on the dorsal surface of the head in the other), but on all other individuals parasitized by both species, members of the two leech species were each found segregated on different body regions (ESM Table 2).

### Patterns of Infestation across Elephant Island

Across all Elephant Island stations, we inspected a total of 68 *C. aceratus*, 709 *C. gunnari,* and 13 *C. rastrospinosus* for the presence of leeches, generally finding relatively low prevalence of leech infestations (Table 3). For infested individuals, each fish species had a unique dominant leech; *T. capitis* was the commonly encountered leech on *C. aceratus* (75%) while *T. bacilliformis* was the most common leech on *C. gunnari* (87.5%). *C. rastrospinosus was infested primarily by N. sawyeri* (50%) (Fig. 2A). Infested individuals displayed a heterogeneous spatial distribution, with most leeches found to the Southwest of the Island (Fig. 2B). Further, leech density was not correlated with fish abundance across stations (Table 4). Despite spatial heterogeneity, there was a clear trend of leeches preferentially infesting only larger adult fishes of all three species [*C. aceratus*: *R*=67–43cm total length (TL), *p*=0.067; *C. gunnari*: *R*=49–34cm TL, *p*=0.000; *C. rastrospinosus*: *R*=56–34cm TL, *p*=0.002; Fig. 2C). Infestation intensity varied by leech species and host fish species. In *C. gunnari, T. bacilliformis* infestations ranged from few individuals to upwards of 20, with a remarkable individual covered by more than 40 leeches. In contrast, *T. capitis* always occurred in comparatively low (<10 individuals) intensities (Fig. 2D).

**Table 3:** Prevalence of infestation of each leech species on each host fish species in the vicinity of Elephant Island and the South Orkney Islands.

| Species | *Elephant Island* |  |  | *South Orkneys* |  |  |
| --- | --- | --- | --- | --- | --- | --- |
|  | *Trulliobdella bacilliformis* | *Trulliobdella capitis* | *Nototheniobdella sawyeri* | *Trulliobdella bacilliformis* | *Trulliobdella capitis* | *Nototheniobdella sawyeri* |
|  | Number infested / Total catch (% prevalence) | Number infested / Total catch (% prevalence) | Number infested / Total catch (% prevalence) | Number infested / Total catch (% prevalence) | Number infested / Total catch (% prevalence) | Number infested / Total catch (% prevalence) |
| *Chaenocephalus aceratus* | 4 / 68 (5.88 %) | 12 / 68 (17.65 %) | 0 / 68 (0%) | 8 / 234 (3.42 %) | 8 / 234 (3.42 %) | 0 / 234 (0 %) |
| *Champscephalus gunnari* | 14 / 709 (1.97 %) | 1 / 709 (0.14 %) | 1 / 709 (0.14 %) | 25 / 333 (7.51 %) | 8 / 333 (2.40 %) | 0 / 333 (0 %) |
| *Chionodraco rastrorpinosus* | 3 / 13 (23.08 %) | 1 / 13 (7.69 %) | 4 / 13 (30.77%) | 1 / 104 (0.96 %) | 7 / 104 (6.73 %) | 0 / 104 (0 %) |

**Figure 2:**
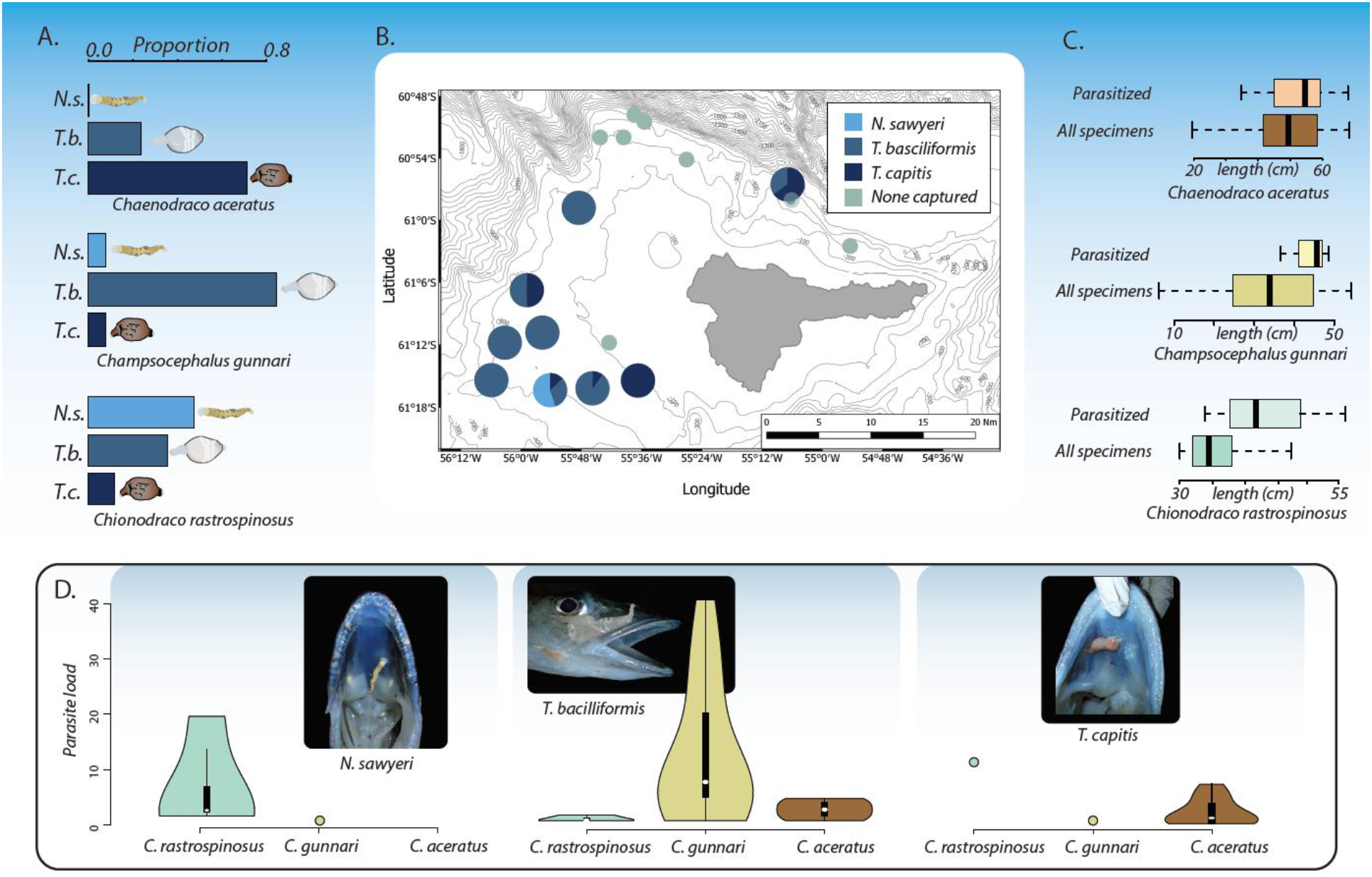
Summary of Antarctic fish leech infestation parameters around Elephant Island. A. Relative proportions of leech taxa found across species of Crocodile icefish. B. Map of stations with pie charts indicating frequencies of leech species found per station. Shadings correspond to leech species. C. Comparison of body size (TL) frequencies of total parasitized individuals to all crocodile icefishes captured. D. Violin plots depicting the infestation intensities by leech species. Embedded boxplots represent the quartiles of the parasite intensities per host-parasite species pair.

**Table 4:**
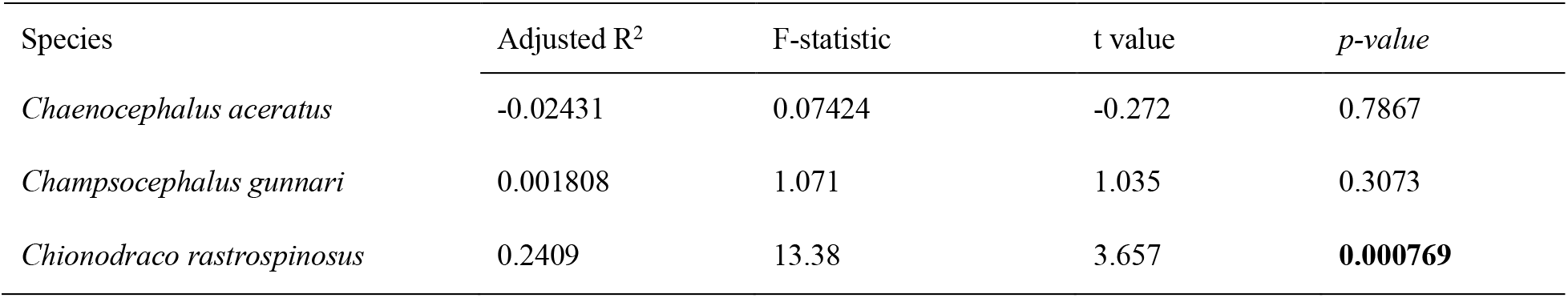
Results of tests for correlation between abundance of parasitized fish and total abundance of fish for each species across stations (at both Elephant Island and South Orkney Islands).

| Species | Adjusted R<sup>2</sup> | F-statistic | t value | p-value |
| --- | --- | --- | --- | --- |
| *Chaenocephalus aceratus* | -0.02431 | 0.07424 | -0.272 | 0.7867 |
| *Champscephalus gunnari* | 0.001808 | 1.071 | 1.035 | 0.3073 |
| *Chionodraco rastrospinosus* | 0.2409 | 13.38 | 3.657 | **0.000769** |

### Patterns of Infestation across the South Orkney Islands

Across the South Orkney Island stations, we inspected a total of 234 *C. aceratus*, 333 *C. gunnari*, and 104 *C. rastrospinosus* for the presence of leeches, and again we observed relatively low prevalence of leech infestations (Table 3). In the South Orkneys, only *T. capitis* and *T. bacilliformis* were encountered on all three fish species. *Trulliobdella capitis* and *T. bacilliformis* were encountered with equal frequency on *C. aceratus* (50% each). Similar to patterns observed at Elephant Island, *T. bacilliformis* was the most common leech on *C. gunnari* (92.6%) (Fig. 3A). Infested individuals again displayed a heterogeneous spatial distribution, with few leeches found in the Eastern portions of the Islands (Fig 3B). Despite spatial heterogeneity, there was a clear trend of leeches preferentially infesting only larger fishes for all three species (*C. aceratus*: *R*=64–38cm TL, *p*=0.018; *C. gunnari*: *R*=52–32cmTL, *p*=0.002; *C. rastrospinosus*: *R*=42–32cm TL, *p*=0.239; Fig 3C). Parasite intensities varied by leech species and host fish species. In *C. aceratus* and *C. gunnari*, *T. bacilliformis* infestations ranged from few individuals up to 14, while only one *T. bacilliformis* individual was observed on a single *C. rastrospinosus*. *Trulliobdella capitis* always occured in low intensities (<5 individuals) on individual fish across all species (Fig. 3D).

**Figure 3:**
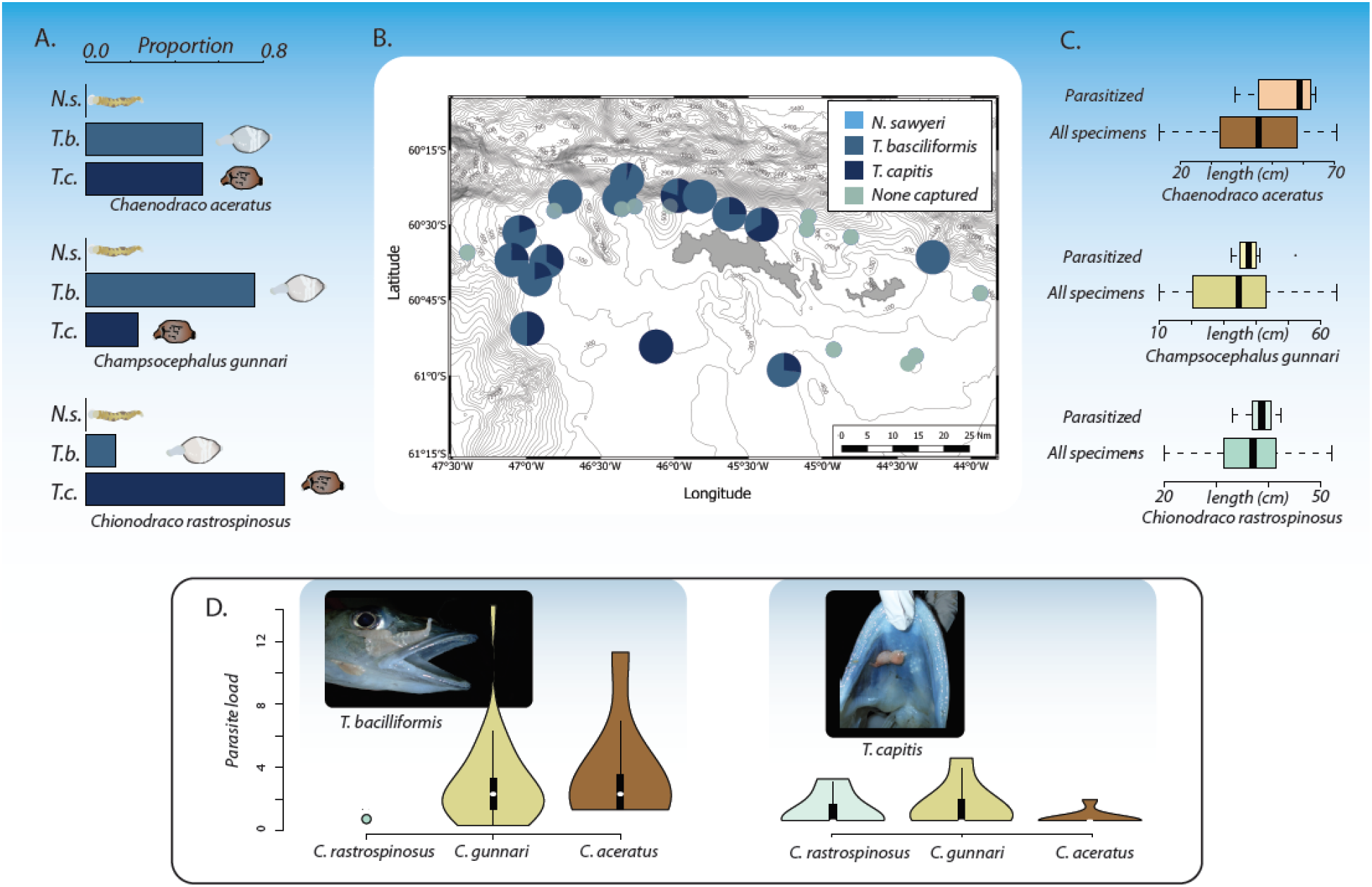
Summary of Antarctic fish leech infestation parameters around the South Orkney Islands. A. Relative proportions of leech taxa found across species of Crocodile icefish. B. Map of stations with pie charts indicating frequencies of leech species found per station. Shadings correspond to leech species. C. Comparison of body size (TL) frequencies of total parasitized fish to all Crocodile icefishes captured. D. Violin plots depicting the infestation intensities by leech species. Embedded boxplots represent the quartiles of the parasite intensities per host-parasite species pair.

## Discussion

Patterns of host-parasite specificity have been commonly observed in marine ecosystems (Adamson and Caira 1994; Whittington et al. 2000; Muñoz and Cortés 2009); however, our results coupled with prior investigations suggest that the Southern Ocean leech species in this study may be better characterized as opportunistic parasites that can utilize a range of host species. While all leech-hosts pairs demonstrated a preference for large-bodied hosts, we did observe asymmetric patterns of parasitism. *Trulliobdella capitis was* observed in relatively high intensities on the host fish species *C. aceratus*, *T. bacilliformis was* encountered in high intensities on the fish species *C. gunnari*, and *N. sawyeri* was observed in high intensity on the fish species *C. rastrospinosus*. Our results further revealed non-random patterns of infestation sites on host bodies. Across all host and parasite species studied, these attachment sites appear restricted to regions of host fishes that exhibit the highest levels of vascularization and thinnest skin. However, attachment site fidelity varied between host-parasite species pairs. This variation suggests two additional drivers of leech parasitism. First, our findings suggest the potential for individual leech species to be dominant at different strata of the water column, yet utilize host fish to passively migrate between strata, and thereby opportunistically encounter additional potential notothenioid hosts. Second, the occurrence of leeches inside the jaws of fish predators suggests the possibility that leeches are passively co-opting the food web to utilize smaller hosts as transmission vectors to reach larger piscivorous icefishes. In total, our findings offer critically needed baseline data for Antarctic fish leech ecology and offer a wide range of novel hypotheses that can be tested as more data on these fascinating organisms becomes available.

### Host-preference in Antarctic leeches?

Previous work has suggested other species of piscicolids to go through boom and bust cycles of population growth, and therefore the parasite infestation prevalence may not be correlated with host abundance (Sawyer and Hammond 1973). Our study supports this hypothesis, demonstrating that leech infestation intensity in Elephant and the South Orkney Islands is not correlated with host fish abundance (Table 4; ESM Tables 3&4). Instead, leech abundance at each island tends to be partitioned between host species. This raises the question of how differences between host fish species in terms of habitat and resource can limit opportunities for between-host transmission that can ultimately promote the evolution of both host-switching and parasite host specificity (Poulin 1992). For example, *C. gunnari* undergoes diel vertical migrations to forage primarily in the water column from close to or in benthic habitats (Kock and Everson 1997). Occurrences of *T. bacilliformis* have been observed on eight other notothenioid species known to feed in the water column, including *Lepidonotothen larseni (Lönnberg* 1905), *Pseudochaenichthys georgianus Norman 1937*, and *Channichthys rhinoceratus Richardson* 1844 (Table 1), suggesting this leech species to be a common parasite for pelagic Antarctic fishes (Utevsky 2007). The occurrence of *T. bacilliformis* infestations on pelagic notothenioids suggests diel vertical migrations expose *C. gunnari* to *T. bacilliformis* at a higher frequency than benthic species. Additionally, *C. gunnari* is also commonly infected by *T. capitis* along the ventral surface of the body, largely between the pelvic fins. *Trulliobdella capitis* has been found on a mixture of pelagic and benthic notothenioids (Table 1), however, there is not enough data available to determine if these leeches are encountered in benthic habitats during vertical migrations of fish hosts, or if these are water column generalists. Given the predominantly ventral attachment, we favor the former hypothesis, however more investigations are needed.

Our results demonstrate that infestation of *C. aceratus* by both species of *Trulliobdella* occurs at highest frequencies on the dorsal surface of the head and inner region of the upper jaw. Several previous studies of the diet of notothenioid fishes have shown that *C. gunnari* is commonly eaten by *C. aceratus* (Flores et al, 2004) - in some cases representing the greatest proportion of the prey items observed in stomach content analysis (Reid et al. 2007). This suggests that predation on *C. gunnari* by *C. aceratus* provides both species of *Trulliobdella* with a transmission pathway to an alternate host. Such a hypothesis would be consistent with the high frequency of parasites found inside the upper jar of this piscivore. This mode of transmission is unusual, as parasite transmission through food webs is often associated with parasites co-opting intermediate hosts as transmission vectors to definitive hosts when the density of the latter is lower than the former (Choisy et al. 2003). However, this is unlikely to be the case here. While the inclusion of intermediate hosts are common for a wide-range of both terrestrial and aquatic parasites with complex life-cycles (Dornburg et al. 2019; Brown et al. 2001; Iglesias-Piñeiro et al. 2016), all leeches have a direct life cycle. Although it is still possible that benthic predators act as aggregators and potential mating sites of both *Trulliobdella* species, it is also possible that *Trulliobdella species* can opportunistically expand their host range. Our finding of *T. bacilliformis* on *C. rastrospinosus* favors a hypothesis of opportunistic host colonization. *Chionodraco rastrospinosus* feeds primarily on krill (*Euphausia superba Dana 1850*), with other fish representing a very small percentage of their diet (Takahashi 1983), making transmission of *T. bacilliformis* by predation less likely.

In addition to host selection, apparent site fidelity of leech species to particular body regions on their fish hosts represents an additional axis of leech ecology that has received little attention in the literature. The results of our study demonstrate that each leech species preferentially infests different body regions on different host fish species. This pattern was not random, and portions of the host body that exhibit low levels of vascularization and increased skin thickness were conspicuously absent from the list of infested locations. Skin thickness has previously been invoked as an explanation for attachment site choice in leeches that parasitize salamanders, as thinner skin provides easier access to a host’s vascular network (Pough 1971). Additionally, areas of the host body with dense and shallow vascular networks should be preferred as these would naturally represent the highest ratio of energy return for investment in foraging effort for sanguivorous feeders. Our results fit both of these criteria. Skin in notothenioids has been found to be thinner in the cranium relative to the post-cranium, with skin in the oral cavity being particularly thin (Eastman 1991). Correspondingly, the cranium is also highly vascularized relative to the trunk (Eastman 1991; Egginton and Rankin 1998), with an additional array of blood vessels providing blood to the pelvic fins (Eastman 1991), possibly to keep the fins extended during perching.

The hypothesis that skin thickness and density of the underlying vascular network drive patterns of site attachment is also supported by prior studies documenting areas of leech infestation on crocodile fishes (Brinkmann 1948; Bielecki et al. 2008; Santoro et al. 2014; Table 1), all of which reported predominantly external areas of the cranium as attachment sides. While leech attachments are temporary, we found high levels of scarring indicative of prior leech attachment on the surfaces of the cranium in numerous individuals (Fig. 4). This provides evidence that the same surfaces are repeatedly used as feeding sites during the lifespan of individual fishes. Previous studies have also included the gill cavity as an attachment point; however, despite investigation of 1,461 individual fishes, we did not observe leeches on the gills or in the gill cavity of any host species. However, additional findings of leeches on gills or in the gill cavity would only further support vascularization and skin thickness of attachment sites to be a limiting factor in the attachment ability of Antarctic marine leeches.

**Figure 4:**
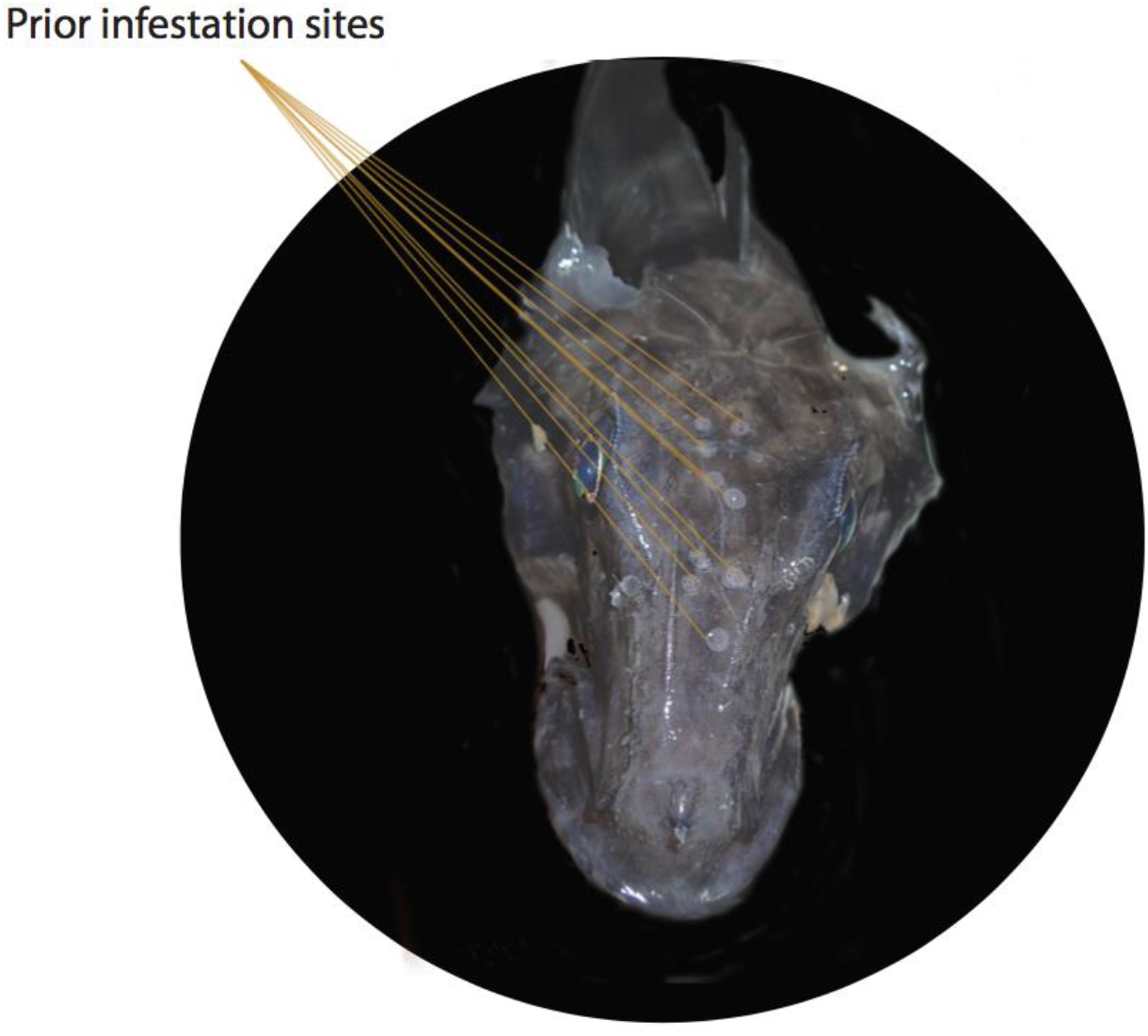
Example of scarring from prior leech infestations on an icefish encountered during the course of this study.

Additionally, our results show that when leeches do occur on the same host, they may be partitioning attachment points in a manner analogous to sympatric species partitioning a landscape (Holmes 1961; Ramasamy et al. 1985; Fuselier and Edds 1994; Arlettaz 1999; Poulin 2001; Humphries et al. 2016). This is especially evident in two *C. aceratus* and four *C. gunnari* individuals that were parasitized by both *T. capitis* and *T. bacilliformis*. In only one individual of *C. gunnari* were the two leech species found to have infected the same body region (one individual of *T. capitis* and one of *T. bacilliformis* between the pelvic fins). Otherwise, whether or not they are observed on the same individual host fish, leech species occupy different body regions of their host, suggesting that leech species may prefer to infest particular body regions. This preference may be a result of variation in host vascularization and skin thickness and how this relates to the ability of the leech to attach to particular body regions or be indicative of competitive interactions (Bashey 2015). However, the low prevalence of leech infestations renders sample sizes too small for further testing expectations of competitive interactions.

## Conclusion

Understanding the relationship between hosts and parasites represents an important, but often neglected, axis when considering forecasts of marine biodiversity under the looming threat of continued environmental change and increased human traffic. Our results provide new baseline data into the ecology and interactions of three species of Antarctic piscicolids with their host fish species. We show these parasites to be highly clustered spatially, but not in a manner that correlates with fish abundance. Instead, we demonstrate low levels of infestation, with high levels of fidelity to select host attachment areas, as well as the potential for trophic transmission between host species. These results reveal an under-appreciated complexity of interactions between this community of hosts and parasites, and offer exciting avenues of further investigations that can illuminate the factors that underlie the abundance and persistence of Antarctic leech parasites. Not only will such studies potentially further our understanding of host-parasite evolution, but also will be essential for contextualizing future changes of Antarctic host-parasite interactions.

## Supporting information

Parker_et_al2019_ESM

## Acknowledgements

We thank the captain and crew of the R/V Cabo de Hornos for their excellent company and assistance during the course of this fieldwork. Also, thanks to Deris S.A. for providing the ship to carry out the work at sea, and to Mr. Enrique Gutiérrez F., general manager of said company, for his permanent support in materializing these tasks. We additionally thank F. Reyda and A. Bielecki for providing helpful reviews of a prior version of this manuscript and an anonymous reviewer for further helpful reviews and proposing the hypothesis that skin thickness and vascularization are driving the site-attachment patterns we observed.

## Compliance with Ethical Standards

The authors declare that they have no competing interests. The authors have followed all applicable national and institutional guidelines for the collection, care, and ethical use of research organisms and material in the conduct of the research, in strict accordance with NOAA Antarctic Ecosystem Research Division policies under ACA Permit #2017-012.

## References

Adamson ML, Caira JN (1994) Evolutionary factors influencing the nature of parasite specificity. Parasitology 109 Suppl:S85–95

Adler D (2005) Vioplot: Violin plot. R package

Arlettaz R (1999) Habitat selection as a major resource partitioning mechanism between the two sympatric sibling bat species *Myotis myotis* and *Myotis blythii*. J Anim Ecol 68:460–471

Bashey F (2015) Within-host competitive interactions as a mechanism for the maintenance of parasite diversity. Philos Trans R Soc Lond B Biol Sci 370.: doi: 10.1098/rstb.2014.0301

Bielecki A, Rokicka M, Ropelewska E, Dziekońska-Rynko J (2008) Leeches (Hirudinida: Piscicolidae)--parasites of Antarctic fish from Channichthyidae family. Wiad Parazytol 54:345–348

Brinkmann A Jr (1948) Some new and remarkable leeches from the Antarctic seas. Sci Results I Kommisjon Hos Jacob Kybwad, Oslo Norw Antarctic Exped 1927-1928 29:1–20

Brown SP, Renaud F, -F. Guégan J, Thomas F (2001) Evolution of trophic transmission in parasites: the need to reach a mating place? J Evol Biol 14:815–820

Bush AO, Lafferty KD, Lotz JM, Shostak W. 1997. Parasitology meets ecology on its own terms: Margolis et al. revisited. Journal of Parasitology 83: 575–583.

Choisy M, Brown SP, Lafferty KD, Thomas F (2003) Evolution of trophic transmission in parasites: why add intermediate hosts? Am Nat 162:172–181

Cocca E, Ratnayake-Lecamwasam M, Parker SK, et al (1995) Genomic remnants of alpha-globin genes in the hemoglobinless antarctic icefishes. Proc Natl Acad Sci U S A 92:1817–1821

Collins MA, Brickle P, Brown J, Belchier M (2010) The Patagonian toothfish: biology, ecology and fishery. Adv Mar Biol 58:227–300

Cruz-Lacierda ER, Toledo JD, Tan-Fermin JD, Burreson EM (2000) Marine leech (Zeylanicobdella arugamensis) infestation in cultured orange-spotted grouper, *Epinephelus coioides*. Aquaculture 185:191–196

De la Mare WK (1994) Estimating confidence intervals for fish stock abundance estimates from trawl surveys. CCAMLR Science

Delord K, Gasco N, Barbraud C, Weimerskirch H (2009) Multivariate effects on seabird bycatch in the legal Patagonian toothfish longline fishery around Crozet and Kerguelen Islands. Polar Biol 33:367–378

Dornburg A, Lamb AD, Warren D, et al (2019) Are geckos paratenic hosts for Caribbean Island Acanthocephalans? Evidence from Gonatodes antillensis and a global review of squamate reptiles acting as transport hosts. Bulletin of the Peabody Museum of Natural History

Duhamel G, Ozouf-Costaz C, Cattaneo-Berrebi G, Berrebi P (1995) Interpopulation relationships in two species of Antarctic fish *Notothenia rossii* and *Champsocephalus gunnari* from the Kerguelen Islands: an allozyme study. Antarct Sci 7.: doi: 10.1017/s0954102095000496

Eastman JT (2000) Antarctic notothenioid fishes as subjects for research in evolutionary biology. Antarct Sci 12.: doi: 10.1017/s0954102000000341

Eastman, JT, & Hikida, RS (1991) Skin structure and vascularization in the Antarctic notothenioid fish Gymnodraco acuticeps. Journal of morphology, 208 (3), 347–365.

Egginton, S, & Rankin, JC (1998) Vascular adaptations for a low pressure/high flow blood supply to locomotory muscles of Antarctic icefish. In Fishes of Antarctica (pp. 185–195). Springer, Milano.

Everson I (2015) Designation and management of large-scale MPAs drawing on the experiences of CCAMLR. Fish Fish 18:145–159

Everson I, Parkes G, Kock K-H, Boyd IL (1999) Variation in standing stock of the mackerel icefish *Champsocephalus gunnari* at South Georgia. J Appl Ecol 36:591–603

Flores, H. Kock, K-H, Wilhelms, S. and C.D. Jones. 2004. Diet of two icefish species from the Southern Shetland Islands and Elephant Island, *Champsocepalus gunnari* and *Chaenocephalus aceratus*. Polar Biol. 27:119–129.

Fuselier L, Edds D (1994) Habitat Partitioning among Three Sympatric Species of Map Turtles, Genus *Graptemys*. J Herpetol 28:154

Gehman A-LM, Hall RJ, Byers JE (2018) Host and parasite thermal ecology jointly determine the effect of climate warming on epidemic dynamics. Proc Natl Acad Sci U S A 115:744–749

Giraldo C, Boutoute M, Mayzaud P, et al (2016) Lipid dynamics in early life stages of the icefish *Chionodraco hamatus* in the Dumont d’Urville Sea (East Antarctica). Polar Biol 40:313–320

Gutt J, Cummings V, Dayton P, et al (2015) Antarctic Marine Animal Forests: Three-Dimensional Communities in Southern Ocean Ecosystems. In: Marine Animal Forests. pp 1–30

Hintze JL, Nelson RD (1998) Violin Plots: A Box Plot-Density Trace Synergism. Am Stat 52:181

Holmes JC. 1961. Effects of concurrent infections on Hymenolepis diminuta (Cestoda) and Moniliformis dubius (Acanthocephala). I. General effects and comparison with crowding. The Journal of parasitology. 47: 209–16.

Humphries NE, Simpson SJ, Wearmouth VJ, Sims DW (2016) Two’s company, three’s a crowd: fine-scale habitat partitioning by depth among sympatric species of marine mesopredator. Mar Ecol Prog Ser 561:173–187

Iglesias-Piñeiro J, González-Warleta M, Castro-Hermida JA, et al (2016) Transmission of *Calicophoron daubneyi* and *Fasciola hepatica* in Galicia (Spain): Temporal follow-up in the intermediate and definitive hosts. Parasit Vectors 9.: doi: 10.1186/s13071-016-1892-8

Jones, Christopher & Sexton, S.N. & Cosgrove III, R.E.. (1999). Surface areas of seabed within the 500 m isobath for regions within the South Shetland Islands (SUBAREA 48.1). 6. 133–140.

Jones, Christopher. (2000). Revised estimates of seabed areas within the 500 M isobath of the South Orkney Islands (Subarea 48.2) and consequences for standing stock biomass estimates of nine species of finfish. CCAMLR science journal of the Scientific Committee and the Commission for the Conservation of Antarctic Marine Living Resources. 7. 197–204.

Karlsbakk E (2004) A trypanosome of Atlantic cod, *Gadus morhua* L., transmitted by the marine leech *Calliobdella nodulifera* (Malm, 1863) (Piscicolidae). Parasitol Res 93:155–158

Kennicutt MC, Chown SL, Cassano JJ, et al (2015) A roadmap for Antarctic and Southern Ocean science for the next two decades and beyond. Antarct Sci 27:3–18

Khan RA (1984) Simultaneous transmission of a piscine piroplasm and trypanosome by a marine leech. J Wildl Dis 20:339–341

Khan RA (1980) The leech as a vector of a fish piroplasm. Can J Zool 58:1631–1637

Kmentová, N., Mašová, Š., Jirounková, E., Nezhybová, V., & Vanhove, M. P. (2016) Leeches of notothenioid fishes in Antarctica: new species discovery and phylogenetic position. In: Proceedings from the 5th Workshop of the European Centre of IchthyoParasitology; November 28-30, 2016; Prušánky, Czech Republic

Kock K-H (2005) Antarctic icefishes (Channichthyidae): a unique family of fishes. A review, Part I. Polar Biol 28:862–895

Kock K-H (2007) Antarctic Marine Living Resources – exploitation and its management in the Southern Ocean. Antarct Sci 19:231

Kock K-H, Everson I (1997) Biology and ecology of mackerel icefish, *Champsocephalus gunnari*: An Antarctic fish lacking hemoglobin. Comp Biochem Physiol A Physiol 118:1067–1077 Kock K-H (1992) Antarctic Fish and Fisheries. Cambridge University Press

Krüger L, Ramos JA, Xavier JC, et al (2017) Projected distributions of Southern Ocean albatrosses, petrels and fisheries as a consequence of climatic change. Ecography 41:195–208

Kuhn T, Zizka VMA, Münster J, et al (2018) Lighten up the dark: metazoan parasites as indicators for the ecology of Antarctic crocodile icefish (Channichthyidae) from the north-west Antarctic Peninsula. PeerJ 6:e4638

Marancik DP, Dove AD, Camus AC (2012) Experimental infection of yellow stingrays Urobatis jamaicensis with the marine leech Branchellion torpedinis. Dis Aquat Organ 101:51–60

Mattiucci S, Nascetti G (2008) Advances and trends in the molecular systematics of anisakid nematodes, with implications for their evolutionary ecology and host-parasite co-evolutionary processes. Adv Parasitol 66:47–148

May RM (1978) Host-Parasitoid Systems in Patchy Environments: A Phenomenological Model. J Anim Ecol 47:833

Melillo D, Varriale S, Giacomelli S, et al (2015) Evolution of the complement system C3 gene in Antarctic teleosts. Mol Immunol 66:299–309

Muñoz G (2014) Parasites communities in the clingfish *Gobiesox marmoratus* from central Chile. Acta Parasitol 59:108–114

Muñoz G, Cortés Y (2009) Parasite communities of a fish assemblage from the intertidal rocky zone of central Chile: similarity and host specificity between temporal and resident fish. Parasitology 136:1291–1303

Muñoz G, Zamora L (2011) Ontogenetic variation in parasite infracommunities of the clingfish *Sicyases sanguineus* (Pisces: Gobiesocidae). J Parasitol 97:14–19

Near TJ, Dornburg A, Kuhn KL, et al (2012) Ancient climate change, antifreeze, and the evolutionary diversification of Antarctic fishes. Proc Natl Acad Sci U S A 109:3434–3439

Oguz MC, Heckmann RA, Cheng CC, et al (2012) Ecto and endoparasites of some fishes from the Antarctic region. Scientia Parasitologica 13:119–128

Pacala SW, Hassell MP, May RM (1990) Host–parasitoid associations in patchy environments. Nature 344:150–153

Palm HW (2011) Fish Parasites as Biological Indicators in a Changing World: Can We Monitor Environmental Impact and Climate Change? In: Progress in Parasitology. pp 223–250

Pennington M (1985) Estimating the Relative Abundance of Fish from a Series of Trawl Surveys. Biometrics 41:197

Pough, FH, 1971. Leech-repellent property of eastern red-spotted newts, Notophthalmus viridescens. Science, 174: 1144–1146.

Poulin R (1992) Determinants of host-specificity in parasites of freshwater fishes. Int J Parasitol 22:753–758

Poulin, R., 2001. Interactions between species and the structure of helminth communities. Parasitology, 122(S1), pp.S3–S11.

Ramasamy P, Ramalingam K, Hanna RE, Halton DW. 1985. Microhabitats of gill parasites (Monogenea and Copepoda) of teleosts (Scomberoides spp.). International Journal for Parasitology. 15: 385–97.

Raymond B, Lea M-A, Patterson T, et al (2014) Important marine habitat off east Antarctica revealed by two decades of multi-species predator tracking. Ecography 38:121–129

Reid WDK, Clarke S, Collins MA, Belchier M (2007) Distribution and ecology of *Chaenocephalus aceratus* (Channichthyidae) around South Georgia and Shag Rocks (Southern Ocean). Polar Biol 30:1523–1533

Ruud JT (1954) Vertebrates without Erythrocytes and Blood Pigment. Nature 173:848–850

Santoro M, Mattiucci S, Cipriani P, et al (2014) Parasite communities of icefish (*Chionodraco hamatus*) in the Ross Sea (Antarctica): influence of the host sex on the helminth infracommunity structure. PLoS One 9:e88876

Sawyer RT, Hammond DL (1973) Observations on the marine leech *Calliobdella carolinensis* (Hirudinea: Piscicolidae), epizootic on the Atlantic menhaden. Biol Bull 145:373–388

Siddall ME, Desser SS (1993) Cytopathological changes induced by *Haemogregarina myoxocephali in its* fish host and leech vector. J Parasitol 79:297–301

Sidell BD, O’Brien KM (2006) When bad things happen to good fish: the loss of hemoglobin and myoglobin expression in Antarctic icefishes. J Exp Biol 209:1791–1802

Smith WOJr, Ainley DG, Cattaneo-Vietti R (2007) Trophic interactions within the Ross Sea continental shelf ecosystem. Philos Trans R Soc Lond B Biol Sci 362:95–111

Takahashi M (1983) Trophic ecology of demersal fish community north of the South Shetland Islands, with notes on the ecological role of krill. Mem Natl Inst Polar Res 23:183–192

Targett TE (1981) Trophic Ecology and Structure of Coastal Antarctic Fish Communities. Mar Ecol Prog Ser 4:243–263

Thompson RM, Mouritsen KN, Poulin R (2004) Importance of parasites and their life cycle characteristics in determining the structure of a large marine food web. J Anim Ecol 74:77–85

Utevsky A (2007) Antarctic piscicolid leeches. Zoologisches Forschungdmuseum Alexander Koenig, Bonn

Whittington ID, Cribb BW, Hamwood TE, Halliday JA (2000) Host-specificity of monogenean (platyhelminth) parasites: a role for anterior adhesive areas? Int J Parasitol 30:305–320

Williams R, Smolenski AJ, White RWG (1994) Mitochondrial DNA variation of *Champsocephalus gunnari* Lönnberg (Pisces: Channichthyidae) stocks on the Kerguelen Plateau, southern Indian Ocean. Antarct Sci 6.: doi: 10.1017/s0954102094000520

