## Supplementary material for "Infestation dynamics between parasitic Antarctic fish leeches (Piscicolidae) and their crocodile icefish hosts (Channichthyidae)": Parker_et_al2019_ESM

### ELECTRONIC SUPPLEMENTAL MATERIALS for

### ESM Tables

| Species | *Elephant Island* | | | *South Orkneys* | | |
|---|---|---|---|---|---|---|
| | *Trulliobdella bacilliformis- Trulliobdella capitis* | *Trulliobdella capitis – Nototheniobdella sawyeri* | *Nototheniobdella sawyeri – Trulliobdella bacilliformis* | *Trulliobdella bacilliformis- Trulliobdella capitis* | *Trulliobdella capitis – Nototheniobdella sawyeri* | *Nototheniobdella sawyeri – Trulliobdella bacilliformis* |
| | *Number infested / Total catch (% prevalence)* | *Number infested / Total catch (% prevalence)* | *Number infested / Total catch (% prevalence)* | *Number infested / Total catch (% prevalence)* | *Number infested / Total catch (% prevalence)* | *Number infested / Total catch (% prevalence)* |
| *Chaenocephalus aceratus* | 0 / 68 (0 %) | 0 / 68 (0 %) | 0 / 68 (0%) | 2 / 234 (0.008 %) | 0 / 234 (0 %) | 0 / 234 (0 %) |
| *Champsocephalus gunnari* | 0 / 709 (0 %) | 0 / 709 (0 %) | 0 / 709 (0 %) | 5 / 333 (0.012 %) | 0 / 333 (0 %) | 0 / 333 (0 %) |
| *Chionodraco rastrospinosus* | 0 / 13 (0 %) | 0 / 13 (0 %) | 0 / 13 (0 %) | 0 / 104 (0 %) | 0 / 104 (0 %) | 0 / 104 (0 %) |

ESM Table 1. Summary of co-infestation prevalence per parasite species pair and host fish species in the vicinity of Elephant Island and the South Orkney Islands.

| Host Species | Leech Species | Body Region | | | | | | |
|---|---|---|---|---|---|---|---|---|
| | | Lateral Surface Head | Dorsal Surface Head | Upper Jaw Inner | Lower Jaw Outer | Between Pectoral Fins | Pectoral Fins | Ventral Surface Body |
| *Champsocephalus gunnari (n=5)* | *Trulliobdella bacilliformis* | 4 / 5 (80%) | 3 / 5 (60%) | 0 / 5 (0%) | 0 / 5 (0%) | 1 / 5 (20%) | 4 /5 (80%) | 0 / 5 (0%) |
| | *Trulliobdella capitis* | 0 / 5 (0%) | 1 / 5 (20%) | 0 / 5 (0%) | 1 / 5 (20%) | 3 / 5 (60%) | 0 /5 (0%) | 1 / 5 (20%) |
| *Chaenocephalus aceratus (n=2)* | *Trulliobdella bacilliformis* | 0 /2 (0%) | 0 /2 (0%) | 1 /2 (50%) | 0 /2 (0%) | 1 /2 (50%) | 0 /2 (0%) | 0 /2 (0%) |

| | | | | | | | |
|---|---|---|---|---|---|---|---|
| *Trulliobdella capitis* | 0 /2 (0%) | 0 /2 (0%) | 0 /2 (0%) | 1 /2 (50%) | 1 /2 (50%) | 0 /2 (0%) | 0 /2 (0%) |

ESM Table 2. Infestation prevalence per leech species and body region for all individuals of *C. gunnari* and *C. aceratus* simultaneously infected by both leech species.

| Area | Species | Stations Total / present | Abundance (#) | S.E. | Lower 95% CI | Upper 95% CI |
|---|---|---|---|---|---|---|
| Elephant Is. | *T. bacilliformis* | 15 / 7 | 1485430 | 992692 | 358833 | 23662300 |
| Elephant Is. | *T. capitis* | 15 / 4 | 376827 | 260126 | 82202.8 | 6144760 |
| S. Orkney Is | *T. bacilliformis* | 21 / 13 | 4204900 | 1843650 | 1808570 | 15746500 |
| S. Orkney Is | *T. capitis* | 21 / 13 | 1485430 | 992692 | 358833 | 23662300 |

ESM Table 3. Estimated total numbers of two species of leeches found on shelf areas of Elephant Island and the South Orkney Islands during the 2018 austral summer.

| Area | Species | Stations Total / present | Abundance (#) | S.E. | Lower 95% CI | Upper 95% CI |
|---|---|---|---|---|---|---|
| Elephant Is. | *C. aceratus* | 15 / 11 | 361173 | 120546 | 124907 | 597438 |
| Elephant Is. | *C. gunnari* | 15 / 13 | 3075333 | 1006649 | 1102337 | 5048329 |
| Elephant Is. | *C. rastrospinosus* | 15 / 4 | 30521 | 9471 | 11958 | 49084 |
| S. Orkney Is. | *C. aceratus* | 21 / 18 | 3401732 | 642957 | 2141559 | 4661905 |
| S. Orkney Is. | *C. gunnari* | 21 / 19 | 2636137 | 940875 | 792056 | 4480218 |
| S. Orkney Is. | *C. rastrospinosus* | 21 / 11 | 25104 | 9722 | 23297 | 26911 |

ESM Table 4. Estimated total numbers of three species of Crocodile icefishes found on shelf areas of Elephant Island and the South Orkney Islands during the 2018 austral summer.

#### ESM Figures

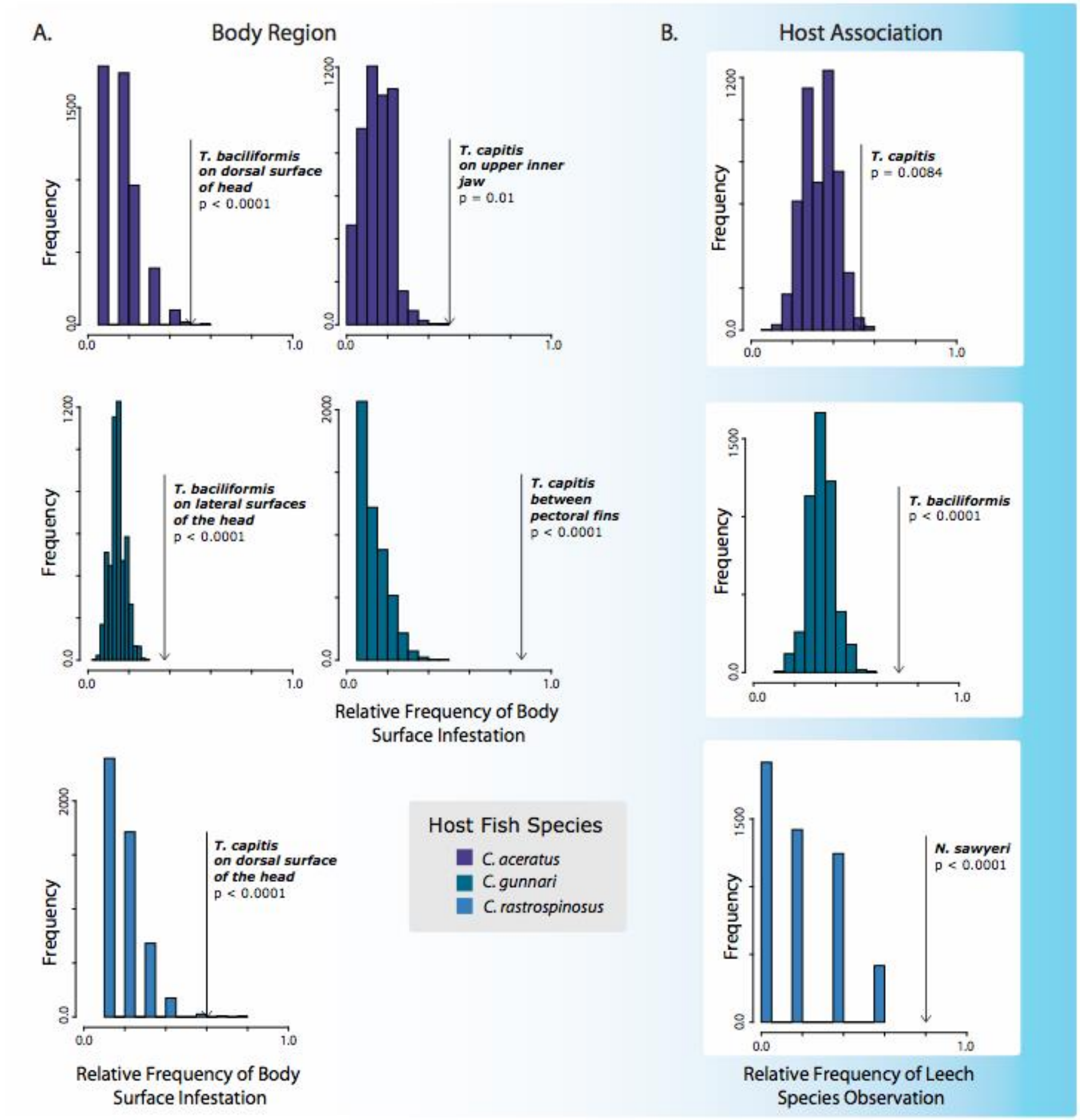


ESM Figure 1: Results of simulations showing (A) expected frequencies of leech species observations on a given body region of a given host fish species under an assumption of random assortment and (B) expected frequency of leech species observation on a given host fish species under an assumption of random assortment. In each histogram, location of arrow represents empirical frequencies of leech species observation calculated in this study.
